# Fully automated workflow for integrated sample digestion and Evotip loading enabling high-throughput clinical proteomics

**DOI:** 10.1101/2023.12.22.573056

**Authors:** Anders H. Kverneland, Florian Harking, Joel Mario Vej-Nielsen, Magnus Huusfeldt, Dorte B. Bekker-Jensen, Inge Marie Svane, Nicolai Bache, Jesper V. Olsen

**Affiliations:** Novo Nordisk Foundation Center for Protein Research, Faculty of Health Sciences, University of Copenhagen, Copenhagen, Denmark; National Center of Cancer Immune Therapy, Department of Oncology, Copenhagen University Hospital - Herlev and Gentofte, Herlev, Denmark; Evosep Biosystems, Odense, Denmark

**Keywords:** Mass spectrometry, workflow automation, plasma proteomics, cancer immune therapy, biomarker discovery

## Abstract

Protein identification and quantification is an important tool for biomarker discovery. With the increased sensitivity and speed of modern mass spectrometers, sample-preparation remains a bottleneck for studying large cohorts. To address this issue, we prepared and evaluated a simple and efficient workflow on the Opentrons OT-2 (OT-2) robot that combines sample digestion, cleanup and Evotip loading in a fully automated manner, allowing the processing of up to 192 samples in 6 hours. Our results demonstrate a highly sensitive workflow yielding both reproducibility and stability even at low sample inputs. The workflow is optimized for minimal sample starting amount to reduce the costs for reagents needed for sample preparation, which is critical when analyzing large biological cohorts. Building on the digesting workflow, we incorporated an automated phosphopeptide enrichment step using magnetic Ti-IMAC beads. This allows for a fully automated proteome and phosphoproteome sample preparation in a single step with high sensitivity. Using the integrated workflow, we evaluated the effects of cancer immune therapy on the plasma proteome in metastatic melanoma patients.

## Introduction

In recent years, mass spectrometry (MS)-based proteomics has become a widely used platform for clinical biomarker discovery and studying cellular signaling networks. This has led to technological developments of fast sequencing mass spectrometers that facilitate short online liquid chromatography tandem mass spectrometry (LC-MS/MS) enabling high-throughput analyses of hundreds of samples per day (1–4). It is increasingly feasible to utilize LC-MS/MS for analyzing large patient cohorts but the bottlenecks for large-scale studies have moved to the pre- and post-analytical sample processing steps, which are unable to keep up with the increased throughput of the LC-MS/MS analyses.

The pre-analytical sample preparation in bottom-up (shotgun) proteomics can crudely be divided into three steps. The first step is preparing the proteins for digestion including protein extraction, cysteine disulfide bond reduction and alkylation. This step typically involves cell lysis or even tissue degradation and will usually be specific to the sample type in question. The second step is the protein digestion with sequence-specific proteases such as trypsin (5). Sequencing-grade proteases are expensive reagents accounting for most of the cost of proteomics sample preparation, and digestion has traditionally been performed overnight at 37°C to optimize enzyme efficiency. The third and final step is preparing the resulting peptide mixture for mass spectrometric analysis by acidifying, desalting and concentrating the sample. The final process has been simplified by sample loading onto reversed-phase C18-based solid-phase material such as an Evotip (Evosep, Odense, Denmark), directly compatible with LC-MS analysis.

While proteome analysis is the cornerstone of LC-MS proteomics, the increased sensitivity and improvements of methodology has also enabled studying post-translational modifications (PTMs) such as site-specific phosphorylations or acetylation sites (6,7). PTMs are mediators of intracellular signaling regulation and by studying these modifications, deep biological insights into cellular dynamics can be attained. Sample preparation for PTM analysis in bottom-up proteomics described above typically requires a PTM-specific enrichment step prior to LC-MS analysis (8,9).

An obvious way to increase the pre-analytical throughput is to automate the sample processing and preparation steps. In addition to increased throughput, automation has the potential to reduce cost and decrease the pre-analytical variability, which is essential to minimize for large cohort studies. Semi-automated sample processing workflows for LC-MS/MS-based proteomics have been developed on the Bravo Liquid Handling Platform (Agilent, Santa Clara, CA) that works with micro-chromatographic cartridges that require manual off-board centrifugation steps (10–12). Liquid handling robots integrating magnetic bead-based sample preparation, such as the Kingfisher Flex (Thermo Scientific, Waltham, MA), enables full automation of the digestion process, but requires relatively large sample input and still involves manual preparation steps before LC-MS analysis (13)

Here, we present a completely automated end-to-end proteomics sample preparation workflow on the Opentrons OT-2 liquid handling robot. The workflow enables simultaneous preparation of up to 192 samples and encompasses the entire process starting from a cell lysate or protein extract to peptide digests loaded on Evotips ready for LC-MS/MS analysis. The workflow is based on magnetic bead protein-aggregation capture (PAC) digestion and can be performed in 6 hours and with an option to include magnetic IMAC-based phosphopeptide enrichment. Through integration of the digestion process with the Evotip loading, this automation strategy enables protein digestion and loading of almost the entire resulting peptide sample, which greatly increases efficiency while reducing the cost of processing.

## Method

### Sample preparation

HeLa lysates were prepared from cells cultured in DMEM media and harvested in boiling 5% sodium dodecyl sulfate (SDS) buffer supplemented with 5 mM tris(2-carboxyethyl)phosphine (TCEP) and 10 mM 2-chloroacetamide (CAA) and incubated for 10 minutes at 99c. Protein concentration was measured using a BCA assay.

Plasma samples for optimization were collected from anonymous healthy individuals. The blood was drawn into sodium citrate 3.2 % tubes (Vacuette, cat# 455322, Greiner BioOne, Kremsmünster, Austria) using a butterfly (Blood Collection Set + Holder 21G × 3/4″, Greiner One Bio, cat# 450085) and immediately centrifuged after collection at 2000 g for 10 min followed by another centrifugation of the supernatant at 3000 g for 10 min to achieve platelet-poor plasma (PPP) and stored at -80°C until analysis.

Patients were recruited and the plasma samples were collected at the Department of Oncology at Herlev Hospital. All enrolled patients provided oral and written informed consent before inclusion and study was approved by the local Ethics Committee (H-15007985). Blood samples were collected at baseline before therapy and after the 1st series of therapy approximately 21 days after. The blood samples were collected in 9 ml K2-EDTA-tubes (Vacuette, cat# 455045, Greiner BioOne) and the plasma was collected after centrifugation at 1300g for 10 min within 2 hours of sample collection and stored at -80c until analysis.

The protein concentration in plasma was approximated to 60 ug/ul. After thawing, the plasma samples were diluted to 225X with PBS and then lysed, reduced and alkylated with 15 min incubation at 37c in 1% SDS, 5 mM TCEP, 10 mM CAA for a final dilution of 300X.

HeLa cell lines for the phosphoproteomics implementation were cultured in DMEM media and seeded into either 96 well or 48 well plates at amounts of 10, 20, 40 or 60 thousand cells per well. Cells were harvested by first washing each well twice in PBS without Magnesium and Calcium. Boiling 1% SDS buffer supplemented with 10 mM TCEP and 20 mM CAA as well as 5 mM β-glycerophosphate, 5 mM sodium fluoride, 1 mM sodium orthovanadate was added and the lysate was incubated at 80°C for 10 min. Lysates were kept in cell culture plates and frozen at -80°C until further processing. Lysates were thawed at 60°C, then treated with 20U Benzonase per well for 10 min at 37°C. Plates were spun down and lysates were transferred into 96 well PCR plates for further processing in the OT-2.

For drug treatment, 40,000 cells were seeded in 48-well plates and grown for 48 hours before treatment with 1uM anisomycin (CAS No. 22862-76-6, A9789-5MG, Sigma Alrdrich) for 0, 0.5, 1 and 2 hours. The samples were analyzed using a 60-SPD gradient on a 15 cm EV1109 column with a steel emitter.

### OT-2 preparation for sample digestion and Evotip loading

The automated PAC workflow is prepared by mixing sample, magnetic beads and acetonitrile in wells of a 96-well sample plate (Eppendorf twin.tec® PCR Plate 96 LoBind®, cat# 0030129512). The samples were diluted until the desired protein input could be contained in 5 ul and then transferred to each well in the sample plate. Immediately before starting the protocol on the OT-2, 5 uL Magnetic beads (MagReSyn hydroxyl beads, cat# MR-HYX2L, ReSyn Biosciences, Ltd, South Africa) and 40 uL MS-grade acetonitrile (ACN) was added to the well for a final aggregation volume of 50 uL.

The OT-2 deck positions including the reservoir plate was set up according to the Evosep step-by-step guide available at https://www.evosep.com/support/automation-opentrons-ot2 with an enzyme:protein mass ratio of 1:25 for trypsin (Cat# T6567, Sigma Aldrich) and 1:100 for endoproteinase Lys-C (Cat# 129-02541, FUJIFILM Wako Pure Chemical Corporation, Richmond, VA) prepared in 50mM Triethyl ammonium bicarbonate (TEAB). Digestion was carried out at room temperature using 4 hours incubation time unless otherwise specified.

Run protocols directly compatible with the Opentrons app was downloaded from the Evosep website where they are available in an easy-to-use HTML format: https://www.evosep.com/support/automation-opentrons-ot2

For the phospho-enrichment protocol, 10 uL Ti-IMAC beads (MagReSyn Ti-IMAC HP, cat# MR-TIM010, ReSyn Biosciences) were pre-prepared in 100% Acetonitrile and placed on the right half of the 96-well plate and 30 uL of lysate mixed with 65 uL ACN and 5 uL Hydroxyl beads (MagReSyn) was placed in the left half of the plate. A solvent reservoir plate was prepared containing ACN (100% v/v), EtOH (100% v/v), Loading buffer (80% ACN, 5% Trifluoroacetic acid (TFA), 1M Glycolic acid, aq.), Wash buffer 2 (70% ACN, 1% TFA, aq.), Wash buffer 3 (20% ACN, 0.1% TFA, aq.), Elution buffer (1% Ammonium, aq.), Digestion buffer (100 mM TEAB), Isopropanol (100% v/v), Enzyme stocks (Trypsin at 1:40 and LysC at 1:80 Protease to protein ratio; stored in acidic conditions before dilution for the digest) and Evotip loading buffer (0.1% FA aq.). A peptide plate was prepared containing 5 uL of (10% v/v) TFA. For running the phospho protocol, a python script was prepared in jupyter notebook (version 6.48).

After protocol run completion, the Evotips were manually moved to the Evosep for LC-MS/MS analysis.

### LC-MS/MS analysis

All samples were eluted online using an Evosep One system (Evosep Biosystems) and separated using an 8 cm (EV1109, Evosep) or 15 cm (EV1137, Evosep) Evosep performance column connected to a steel emitter (EV1086, Evosep) and heated to 40°C. The 100 SPD, 60 SPD or 30 SPD methods were used.

The eluted peptides were analyzed on an Orbitrap Exploris Mass Spectrometer (Thermo Fisher Scientific, Waltham, MA) applying 2 kV spray voltage, funnel RF level at 40, and a heated capillary temperature set to 275°C. The mass spectrometer was operated in positive mode using data independent acquisition (DIA).

In DIA mode, full scan spectra precursor spectra (350–1400 Da) were acquired with a resolution of 120,000 at m/z 200, a normalized AGC target of 300%, and a maximum injection time of 45 ms. Fragment spectra were recorded in profile mode fragmenting 49 consecutive 13 Da windows (1 m/z overlap) covering the mass range 361–1033 Da with a resolution of 15000. Isolated precursors were fragmented in the HCD cell using 27% normalized collision energy, a normalized AGC target of 1000%, and a maximum injection time of 22 ms.

For phosphoproteomics acquisition, full scan spectra precursor spectra (472–1143 Da) were acquired with a resolution of 120,000 at m/z 200, a normalized AGC target of 300%, and a maximum injection time of 45 ms. Fragment spectra were recorded in profile mode fragmenting 17 consecutive 39.5 Da windows (1 m/z overlap) covering the mass range 472– 1143 Da with a resolution of 45,000. Isolated precursors were fragmented in the HCD cell using 27% normalized collision energy, a normalized AGC target of 1000%, and a maximum injection time of 86 ms.

### Data analysis

The MS RAW-files were analyzed using Spectronaut v18 (Biognosys AG, Schlieren, Switzerland) with directDIA+ using default search settings against a FASTA file containing the human proteome (SwissProt, 20,431 sequences, with signal peptides removed) and the sequences of the two proteolytic enzymes. For the patient samples an additional FASTA file containing the protein sequences for the checkpoint inhibitor drugs was included.

Carbamidomethyl was set as a fixed modification and N-terminal acetylation and oxidation of methionine as variable modifications. A Q-value of 1% against mutated decoys was applied to filter identifications. Quantification was performed using the automatic setting and normalized with the integrated cross-run normalization feature unless otherwise specified.

The method evaluation feature was applied when comparing different sample preparations within the same search. Phosphoproteomics data additionally was searched using Phosphorylation (STY) as a variable modification in the PTM tab, with a localization probability cutoff of 0.75.

All data analysis was performed in R version 4.2.2 (R Core Team (2021). R: A language and environment for statistical computing. R Foundation for Statistical Computing, Vienna, Austria) with R Studio 2023.09.1 Build 494. The patient data was normalized using the Variance Stabilization Normalization using vsn package (14). For statistical purposes, missing data in the patient samples were imputed using the MissForest package after filtering out proteins found in less than 30% of all samples (15). The outcome groups were compared by comparing the log2 of the ratio after/before therapy using a linear model from the limma package (16). P-values were adjusted using the Benjamini-Hochberg procedure. The quantified plasma proteins were annotated using an manual adaption of the human secretome found at the human protein atlas (17)

Phosphorylation site stoichiometry information was calculated using a plugin implemented into the Perseus platform (Rapid and site-specific deep phosphoproteome profiling by data-independent acquisition (DIA) without the need for spectral libraries) (18). The exported data was processed using the dapar/Prostar package (19). First, the data was filtered for missingness below 25% in all samples. Data was normalized using the Variance Stabilization Normalisation. Missing values were imputed using the slsa algorithm for partially observed values and determined quantile imputation was done for data points missing in entire conditions (20). Regulated sites were identified using students t test between each condition with log2 fold change cutoff at 0.5. P-values were adjusted using the Benjamini-Hochberg algorithm. Kinase activity estimation was done for regulated sites using the ROKAI tool (21).

## Results

### Sample preparation on the Opentrons OT-2 robot

The fully automated proteomics sample preparation workflow was implemented on the OT-2 to initially process up to 96 samples in parallel within 6 hours. This involved 20 min protein aggregation capture (PAC) on magnetic beads, 10 min buffer exchange and washing, up to 4 hours on-bead protease digestion at room temperature (RT), and finally 60 min Evotip loading (Figure 1A). The OT-2 was chosen, as it is flexible and accessible to most laboratories due to its low cost and straightforward programming control interface. The layout of the OT-2 deck for automated sample preparation of proteome samples includes buffers, Evotips and carefully considered usage of pipetting tips to avoid manual intervention (Figure 1B). Traditionally, tryptic digestion prior to LC-MS/MS analysis has been carried out at 37°C and overnight to ensure complete digestion, which requires either a heating module or implementing a manual step to move the sample plate to a heating device. Instead, avoid manual intervention and maximize throughput, we tested the actual digestion effieciency when combining endoproteinase Lys-C and trypsin digestion at different temperatures and with much shorter digestion periods. Resulting peptide mixtures on Evotips were analyzed by online LC-MS/MS using data-dependent acquisition (DDA) on a Fusion Lumos (Thermo Fisher Scientific) MS instrument (13). We found that the overnight digestion performed equally well at room temperature as at 37°C achieving ∼10% missed-cleavage sites (Figure 1C). At shorter time periods, digestion at 37°C was marginally better than room temperature with the most pronounced effect at the shortest time period of 1 hour. Thus, our experiment demonstrated that short 2-4h digestion periods at room temperature is feasible and only a minor compromise in digestion efficiency.

**Figure 1:**
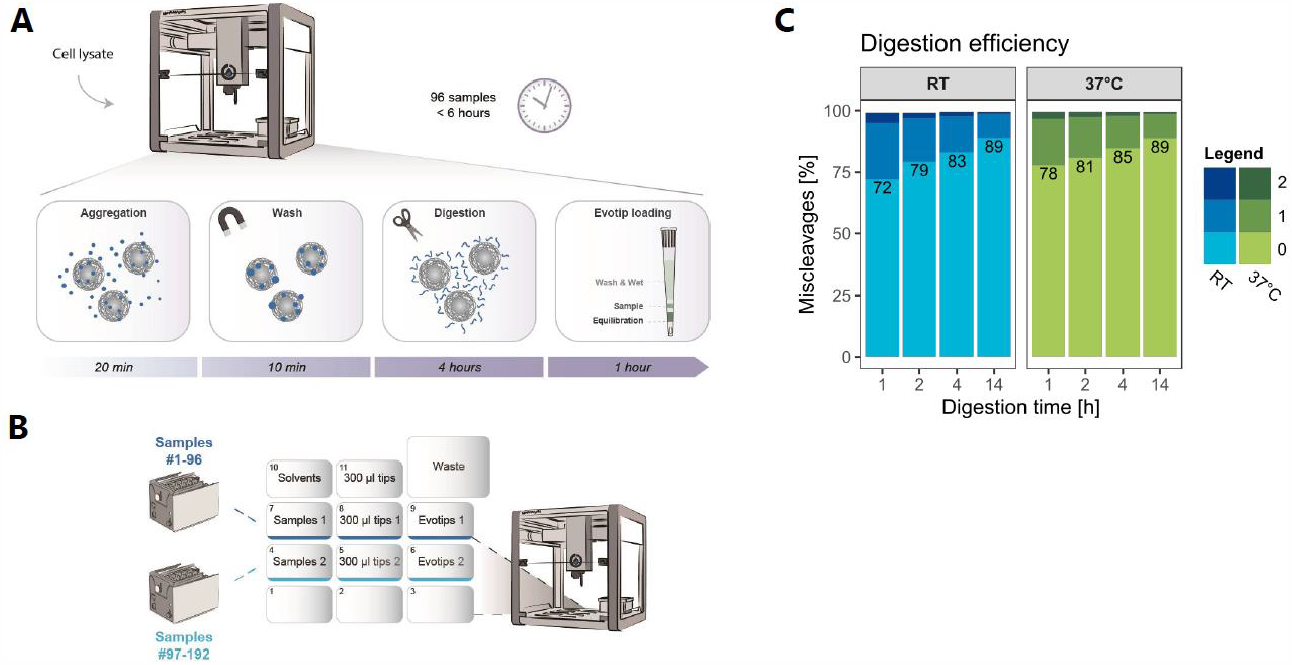
Overview of the automated workflow. **A**: Schematic overview of the integrated workflow on Opentons OT-2 robot. **B**: Deck layout for both 96 and 192 samples of the OT-2 before digestion. **C**: Comparison of digestion efficiency at different temperatures and incubation periods.

### Performance in HeLa cell lines

We started by evaluating the automated workflow using HeLa cell lysate, which is a well-known and frequently used standard in LC-MS/MS-based proteomics quality control. For matching the high-throughput of the automated workflow with LC-MS/MS analysis measurement speed, we decided to test the performance using a 100 samples-per-day (SPD) online LC gradient. Using HeLa cell lysate input equivalent to only 1-ug total protein, we were able to quantify almost 50,000 peptides and 5600 proteins (Figure 2A and 2B). This coverage did not increase when increasing the sample mass input to 15 ug starting material and analyzing a fraction of resulting peptides equivalent to the 1-ug input. These results indicate that sample losses when preparing cell lysates with 1-ug protein are neglectable. To test the performance of the workflow with deeper proteome profiling, we also ran the samples on the considerably longer 30 SPD gradient. The level of quantified peptides and proteins were considerably higher, but still only discretely affected by the sample input. The coefficient of variance (CV) was as expected lower with the longer gradient given the higher coverage (Figure 2C). Similarly, data completeness was also more complete with the 30SPD gradient (Figure 2D). The Pearson correlation coefficients between the workflow replicates were comparable between the gradients demonstrating reproducible protein quantifications (Figure 2E). A list of the quantified proteins and the median quantity can be found in the supplementary table 1.

**Table 1:** Overview of patient cohort:

|  | *All* | *Responders* | *Non-Responders* |
| --- | --- | --- | --- |
| **No. patients, n (%)** | 48 (100) | 26 (54) | 22 (46) |
| **Age, median (min-max)** | 72.5 (27-86) | 71 (44-83) | 73,5 (27-86) |
| **Sex, Male:Female** | 36:12:00 | 21:05 | 15:07 |
| **Melanoma type, n** |  |  |  |
| Skin or unknown focus | 47 | 26 | 21 |
| Ocular | 1 | 0 | 1 |
| **Treatment, n** |  |  |  |
| Pembrolizumab | 32 | 19 | 13 |
| Ipilimumab/Nivolumab | 14 | 7 | 7 |
| Ipilimumab | 2 | 0 | 2 |
| **Best overall response, n** |  |  |  |
| Complete response (CR) | 5 | 5 | 0 |
| Partial response (PR) | 12 | 12 | 0 |
| Stabile disease (SD) | 12 | 9 | 3 |
| Progressive disease (PD) | 19 | 0 | 19 |
| **Follow-up, median days (min-max)** |  |  |  |
| Progression-free survival (PFS) | 233 (33-852) | 478.5 (195-852) | 83.5 (22-176) |
| Overall survival (OS) | 537 (69-858) | 652.5 (395-858) | 314 (69-813) |

**Figure 2:**
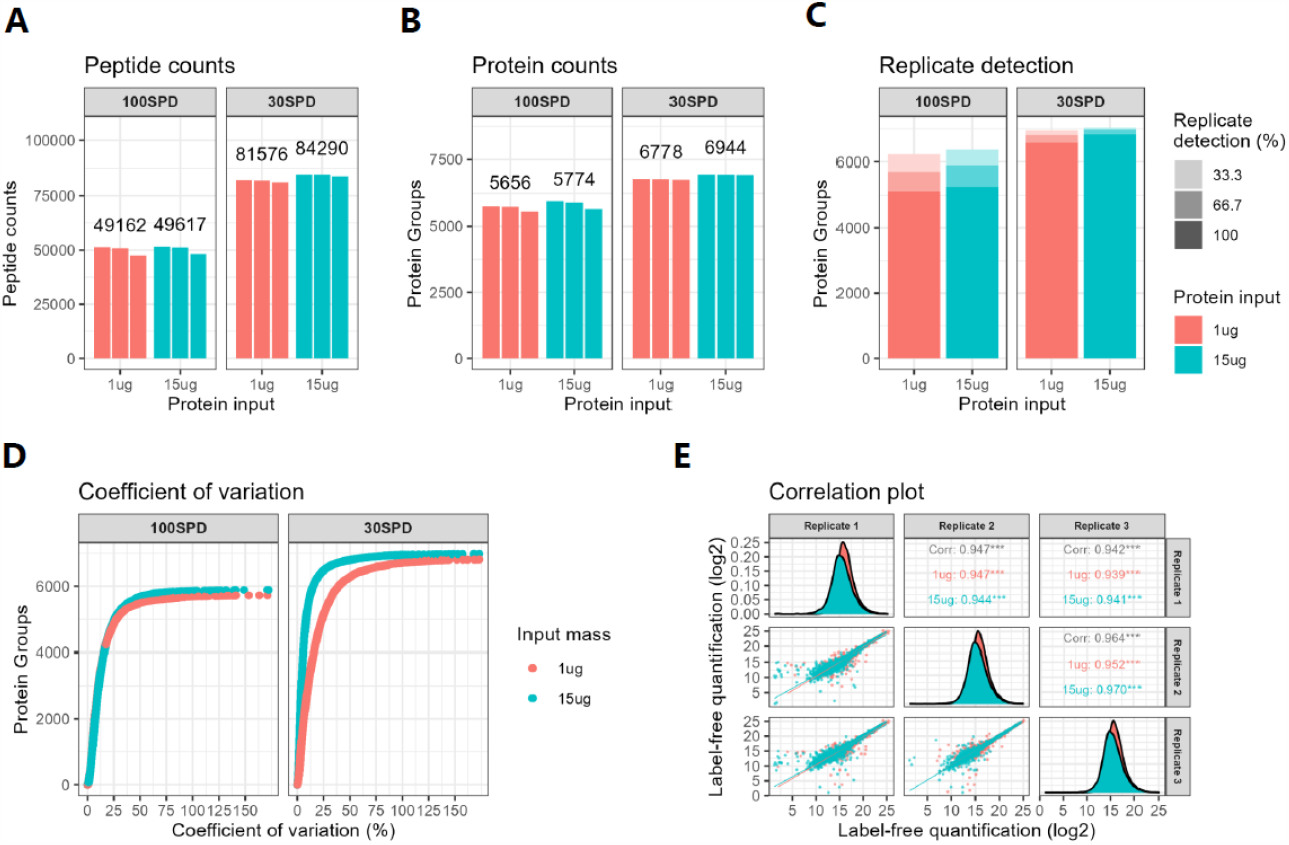
Performance in HeLa samples. **A**: Count of identified peptides in three workflow replicates at 100SPD and 30SPD gradients with 1 ug or 15 ug protein input. **B**: Count of unique protein groups identified in three workflow replicates at 100SPD and 30SPD gradients with 1 ug or 15 ug protein input. **C**: Detection of protein groups across three workflow replicates at 100SPD and 30SPD gradients with 1 ug or 15 ug protein input. **D**: Coefficient of variation comparison of protein inputs and gradient lengths. **E**: Correlation between the workflow replicates with 1 ug input at 100SPD and 15 ug at 30SPD.

We next tested the robustness and reproducibility of the sample preparation by analyzing a large number of workflow replicates back-to-back. When running 32 sample replicates, the performance of the peptide quantification depth was highly comparable (Figure 3A). The CV between the samples was low with approximately 2000 proteins below 10% CV and 3300 below 20% CV (Figure 3B). The CV of the individual protein was as expected correlated to its relative abundance in the sample (Figure 3C).

**Figure 3:**
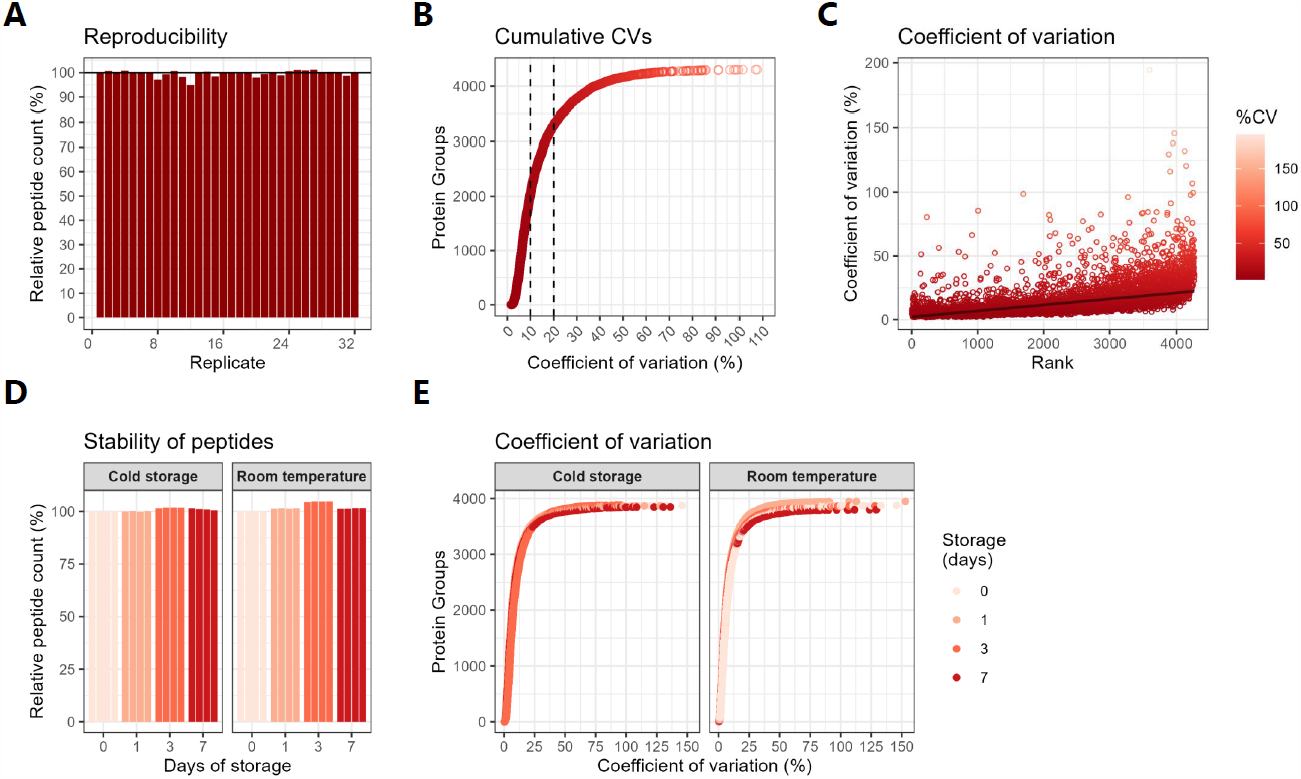
Reproducibility and stability of the automated workflow. **A**: Relative count of identified peptides across 32 workflow replicates. **B**: Cumulative coefficient of variation (CV) across 32 workflow replicates. **C**: Coefficient of variation (CV) plottet against the relative abundance rank of the protein group. **D**: Relative peptide count after up to 7 days of storage either cold or at room temperature. **E**: Coefficient of variation after up to 7 days of storage.

Samples ready for LC-MS/MS cannot always be analyzed immediately after sample preparation. A convenient workflow would therefore allow sample storage until MS measurement becomes available. Since the final step in the automated workflow is sample loading on Evotips, the possibility to store a peptide digest is limited by its stability on the Evotip. We tested the impact of storing samples either cold (4°C) or at room temperature (25°C) up to seven days before the LC-MS injection. The number of quantified peptides and their CVs remained stable through the testing period at both storage conditions (Figure 3D and 3E).

### Performance in plasma samples

Automated sample preparation is a prerequisite for handling and analyzing large-scale clinical cohort studies. Clinical proteomics of large patient cohorts often involves analysis of blood plasma. Plasma is a rich source of proteins and represents systemic physiology. The extreme dynamic range of plasma proteome of ∼12 orders of magnitude is a key challenge for LC-MS/MS based analyses, but plasma remains an attractive source of biomarkers that is routinely collected in most clinical cohorts. We therefore tested the performance of the automated workflow with plasma samples using only 1 ug of protein input using the 100 SPD method. The maximum number of identified peptides and proteins were 3134 and 386, respectively (Figure 4A and 4B), which is comparable to state-of-the-art DIA analysis in undepleted plasma using short LC gradients (1,22,23). Interestingly, the best performance was seen with the lowest sample load of 250 ng and 500 ng corresponding to 25% and 500 ng of the sample input. Likely, due to column saturation of peptides derived from the most abundant plasma proteins at higher sample loads. This trend was also confirmed when comparing the CV and protein sequence coverage of the different sample loads where 250 ng was slightly better than 500 ng (Figure 4C and 4D). We are able quantify proteins within multiple different functions many of which were listed as FDA drug targets (Figure 4E and 4F). A list of the quantified proteins, their annotation and the median quantity can be found in the supplementary table 2.

**Figure 4:**
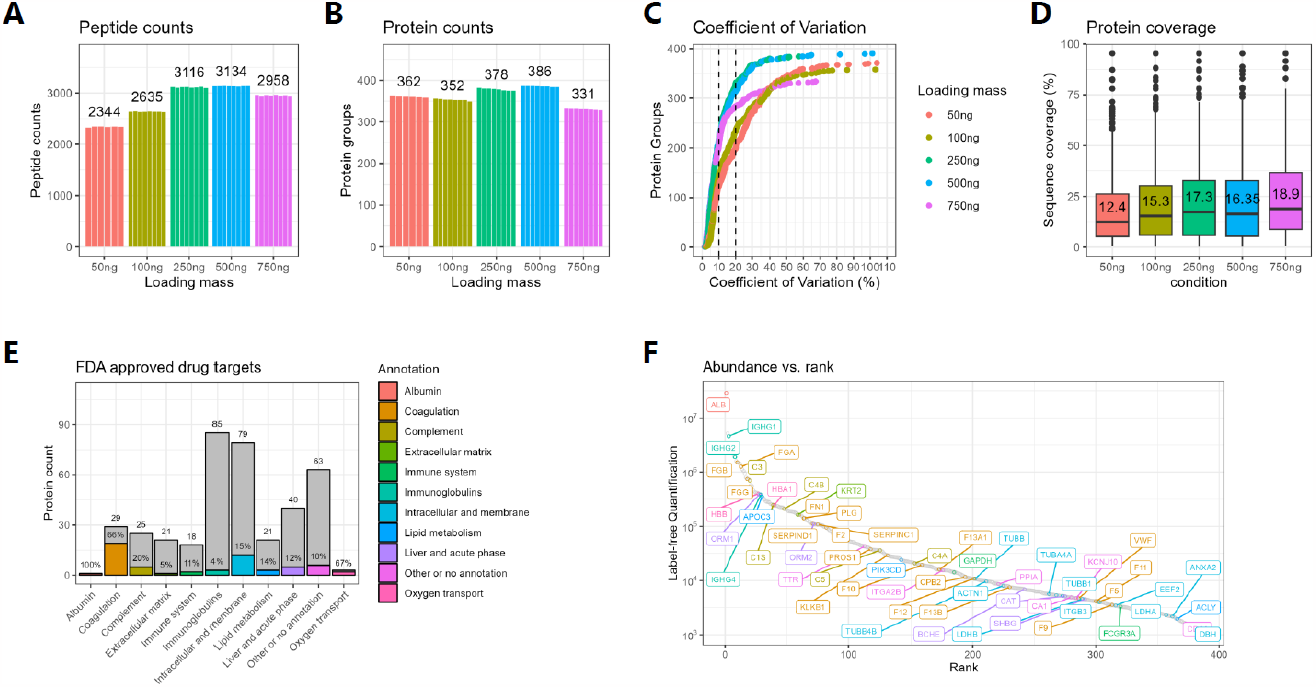
Performance in plasma samples. **A**: Identified peptides with 1 ug plasma using different Evotip loading masses. **B**: Identified protein groups with 1 ug plasma using different Evotip loading masses. **C**: Coefficient of variation (CV) across different loading masses. **D**: Relative protein sequence coverage with different loading masses. **E**: Annotation overview of identified proteins adapted from the proteinatlas.org with the relative number of FDA approved drug targets for the 250ng sample load. **F**: Relative abundance rank vs median label-free intensity (LFQ) in the 250ng load. FDA approved drug targets are labelled and colored according to protein annotation.

### Clinical application in cancer immune therapy

To test the automated workflow in a real-life clinical setting, we analyzed the plasma proteome from a clinical cohort consisting of 48 metastatic malignant melanoma patients with paired plasma samples collected in a biomarker study (Figure 5A). The patient samples were collected before and 3 weeks after initiating of immune therapy with checkpoint inhibitors (CPIs). The outcome of the therapy was assessed with Positron emission tomography–computed tomography (PET-CT) after 3 and 6 months.

**Figure 5:**
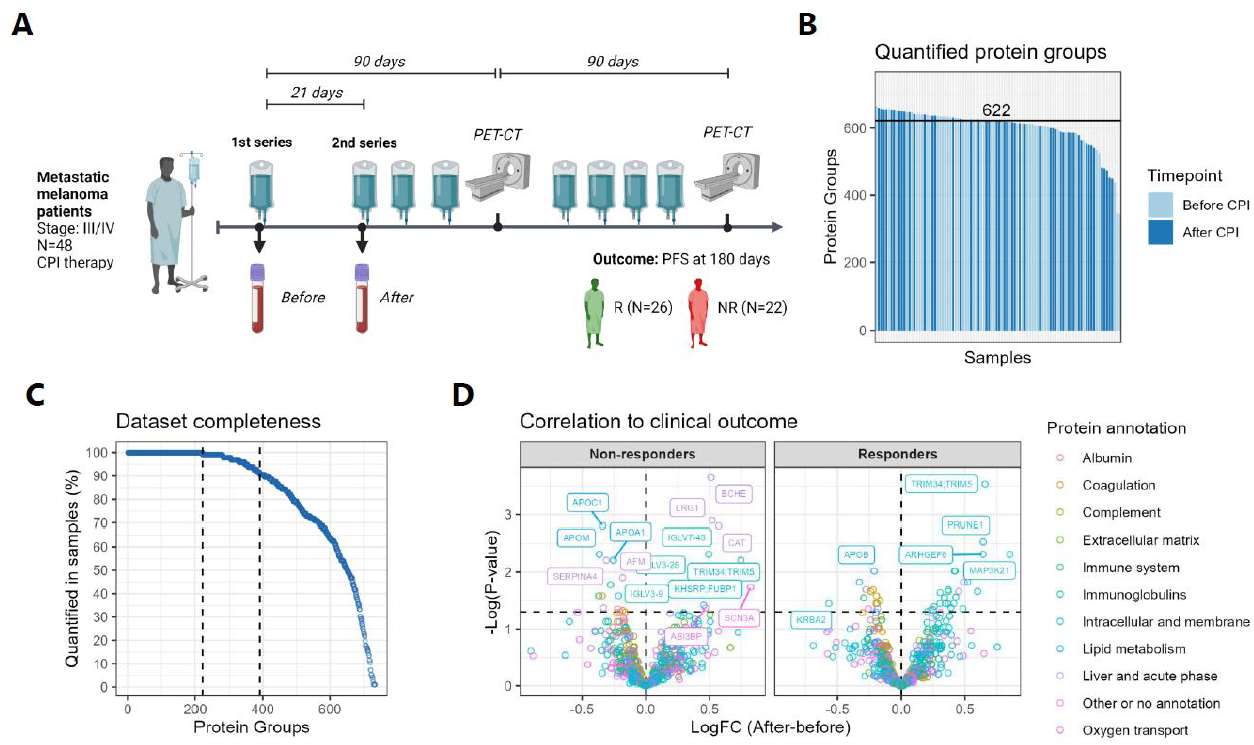
Application in clinical cohort of metastatic melanoma patients. A: Overview of study design, patient inclusion and evaluation. The melanoma patients were recruited prior to checkpoint inhibitor therapy where the first sample was collected (before CPI) and a 2nd blood sample was collected approximately 21 days after the 1st therapy (after CPI). Clinical response to therapy was determined as progression free survival of more than 180 days encompassing two routine evaluation scans. B: Count of quantified proteins across all patients and both time points. C: Dataset completeness assessment visualizing the protein groups identifications across the dataset. D: Volcano plot comparing the time points (after CPI - before CPI) within responding and non-responding patients.

Across all neat plasma samples, we were able to quantify a median of 622 protein groups with approximately 200 proteins quantified in all samples and 400 proteins quantified in 90% of the samples (Figure 5B and 5C). A list of the quantified proteins and the median quantity can be found in the supplementary table 3.

To evaluate the potential for biomarker discovery in this patient cohort, we compared the dynamics of protein levels from before to after initiating of therapy in responding patients and non-responding patients (Figure 5D). These comparisons showed that non-responding patients had higher increases in 3 liver and acute phase proteins (BCHE, CAT, LRG1) and decreases in several apolipoproteins (APOA1, APOC1, APOM). High LRG1 levels have previously been reported as a poor prognostic marker in malignant melanoma and associated with disease relapse in the neoadjuvant setting (24). Similarly decreases of several apolipoproteins (APOB, APOE, APOM) in non-responding patients have also previously been reported as prognostic markers for malignant melanoma (23). Responding patients had increases in several intracellular proteins (ARHGEF6, PRUNE1, MAP4K21, TRIM34;TRIM5). ARHGEF6 has been linked to T-cell migration in lung cancer (25) while MAP3K21 (or MLK4) has also been implicated in immune infiltration in cervical cancer (26,27). This analysis demonstrates the potential for biomarker discovery with clinical proteomics samples using the fully automated sample preparation workflow.

### Phosphoproteomics

Global analysis of site-specific protein phosphorylation status provides a different view of the proteome by directly informing on the activity states of signaling pathways and networks (28). However, as phosphorylation is a sub-stoichiometric modification, it typically requires implementation of specific phosphopeptide enrichment strategies in the sample preparation workflows. To do this, we modified the configuration of the OT-2 protocol to incorporate an automated phosphopeptide-enrichment step after the peptide extraction and digestion. While a small part of the peptide digest was loaded on Evotips for proteome analysis, the remaining sample was enriched for phosphopeptides and subsequently loaded on Evotips for phosphoproteome analysis. The 96-well plate located in the magnetic unit housing the magnetic beads was used for both the proteome digestion and the phosphopeptide enrichment, allowing 48 samples per plate. A peptide plate was prepared for the processing of peptides in between the digest and the phosphopeptide enrichment and acidification of the phosphopeptides after elution in basic ammonia buffer (Fig. 6A).

**Figure 6:**
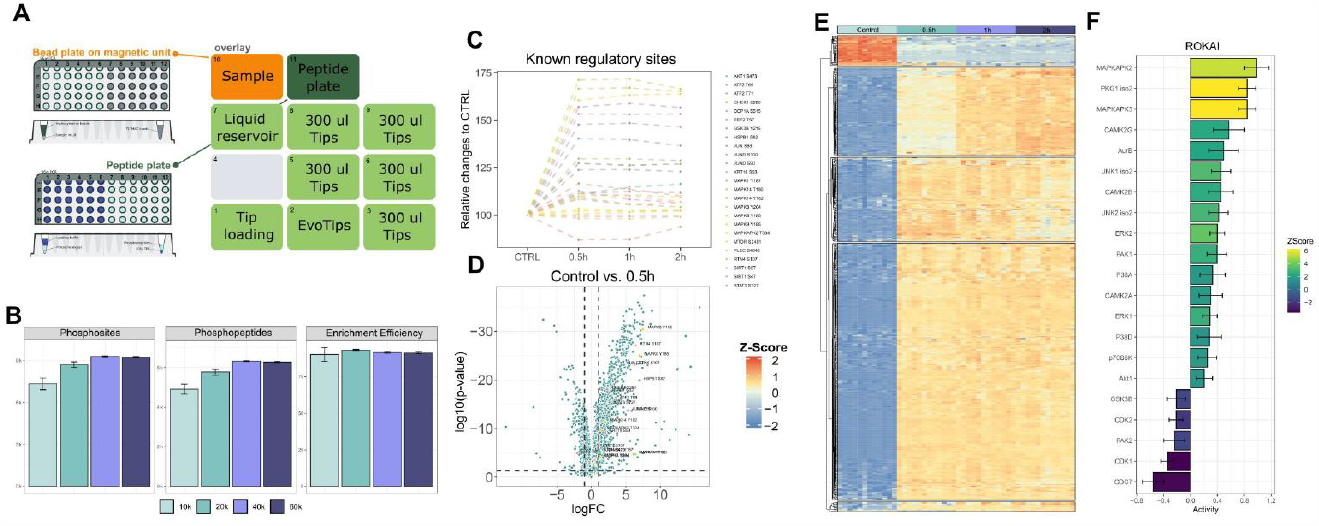
Application of Phosphoproteomics. **A**: Overview on the OT-2 setup for phosphoproteomics. **B**: Initial benchmarking of the using HeLa cell lines. 10,000 and 20,000 cells were seeded in a 96 well format and 40,000 and 60,000 were seeded in a 48 well format. **C**: Relative changes of known interactors to Anisomycin treatment, based on Phosphosite Plus annotated sites. **D**: Volcano plot of the control condition against 0.5h of treatment. Known interactors are labeled. **E**: Heatmap of regulated sites > 2 log FC for the different timepoints. **F**: Rokai plot of regulated sites for the 0.5 h timepoint against control.

To test the performance and sensitivity of this approach, we seeded 10,000 or 20,000 HeLa cells per well in a 96-well plate and 40,000 or 60,000 cells per well in a 48-well plate, respectively. The cells were harvested after 2 days after seeding and subsequently processed with the phosphoproteomics protocol on the OT-2. In all conditions, we were able to identify more than 7,000 phosphosites and 4,000 phosphopeptides, with 40,000 cells in a 48 well plate giving the best results with >8000 phosphosites reproducibly identified. Sample acidification and dilution of the digest directly in the binding buffer was sufficient for phosphopeptide enrichment using Ti-IMAC beads, resulting in enrichment efficiencies of >90% - without any C18 cleanup prior to enrichment (Figure 6B).

To assess the full potential of such an automated phosphoproteomics workflow for high-throughput cell signaling studies, we carried-out a large-scale experiment analyzing dynamic phosphorylation sites in response to anisomycin treatment. Anisomycin is a bacterial antibiotic that inhibits protein synthesis and induces a cellular stress response involving protein kinases such as JNK and p38 (29). After drug incubation for up to 2h, we could see clear temporal phosphorylation dynamics of known anisomycin-regulated sites and many sites were markedly up-regulated already within half an hour of stimulation (Figure 6C). To assess reproducibility of the workflow, we compared the 0.5 h anisomycin treatment time point with the unstimulated control across 12 replicate samples (Figure 6D). We found >1000 significantly regulated sites including up-regulated sites on known downstream targets of JNK and p38. Unsupervised k-means clustering of all regulated sites showed distinct clusters related to the activation or inhibition of phosphosites over time (Figure 6E). A Rokai analysis pinpointed the changes in predicted kinase activity associated to p38 and JNK downstream signaling as expected by the drug activity (Figure 6F).

## Discussion

Fast and reliable sample preparation is becoming increasingly important for bottom-up proteomics with the continuous improvements of sensitivity and throughput of the LC-MS/MS platform. Poor sample preparation results in inaccurate and irreproducible measurements or introduces contaminants that can interfere with the downstream analysis. Even when done stringently, manual sample preparation is laborious and prone to inter-investigator heterogeneity. Especially when the technological advances in mass spectrometric instrumentation enable faster LC-MS/MS measurements, reproducible, scalable and high-throughput sample preparation will be key for successful proteomics application to increasingly larger sample cohorts.

Here we presented a fully automated, hands-off, end-to-end proteomics sample preparation workflow based on the versatile and affordable Opentrons OT-2 platform. The workflow combines the PAC digestion using magnetic micro-beads with automated loading of resulting peptides directly on Evotips for reproducible desalting and storage of samples. The workflow requires no manual intervention after it has been started, resulting in ready-to-analyze samples in less than 6 hours for up to 96 samples in parallel. With immediate capture and storage of peptides on Evotips, the protocol offers an alternative to bulk material preparation by emphasizing efficient utilization of the sample. This greatly reduces the amount and cost of sequencing grade proteases for digestion while preserving the unprocessed sample for later usage. Recently, a parallel effort to automate sample preparation on the OT-2 platform demonstrated equal reproducible performance for LC-MS/MS analysis but at lower throughput including overnight digestion at 37°C and 30 SPD LC gradients (30). Moreover, the script generation in this platform is segmented into several subprocesses providing a very flexible implementation but also requiring some manual intervention. In comparison, the protocol used in this study is designed to be directly compatible with the Opentrons app and can be downloaded from the Evosep website, where they are available in an easy-to-use HTML format.

Our results demonstrated that the workflow is reproducible and stable with a proteome depth on par with what would be expected in manual workflows. In plasma samples, we see that the loading for LC-MS/MS analysis should be reduced for optimal performance on the LC-MS/MS.

When analyzing clinical samples by LC-MS/MS, the biological variability is high, and it is therefore important to minimize variation in the pre-analytical sample handling and preparation. In this study, we demonstrate how an automated workflow can be readily used for biomarker discovery in clinical plasma samples.

The addition of automated phosphopeptide enrichment using magnetic Ti-IMAC beads is integrated and only adds a few additional hours to the total run time. Although the number of samples that can be processed is half of that when preparing proteomes, this workflow allows for up to 48 proteomes and 48 phosphoproteomes in parallel enabling streamlined cell signaling and systems biology investigations.

## Conclusion

Robust and reproducible automation of the sample preparation for bottom-up proteomics and phosphoproteomics is feasible and easily achieved on the Opentrons OT-2 robot. The performance in cancer cell lines and plasma of the automated system is on par with what would be expected in a manual workflow and at higher throughput. Integration of the digestion and sample loading on Evotips allows minimal hands-on time, high throughput and a substantial decrease in the required sample input and reagents needed for digestion. We demonstrate the performance and potential for large-scale biomarker and cellular signaling studies.

## Acknowledgement

Work at The Novo Nordisk Foundation Center for Protein Research (CPR) is funded in part by a donation from the Novo Nordisk Foundation (NNF14CC0001). Part of the work was carried out as a part of the BRIDGE – Translational Excellence Programme funded by the Novo Nordisk Foundation (NNF20SA0064340). The proteomics technology developments applied were part of a project that has received funding from the Innovation Fund Denmark under the grant ERA-PerMed-JTC2022-OVA-PDM and the European Union’s Horizon 2020 research and innovation program under grant agreement EPIC-XS-823839, PUSHH-861389, and ERC synergy grant HighResCells-810057.

